# Ultrasound Produces Extensive Brain Activation via a Cochlear Pathway

**DOI:** 10.1101/233189

**Authors:** Hongsun Guo, Mark Hamilton, Sarah J. Offutt, Cory D. Gloeckner, Tianqi Li, Yohan Kim, Wynn Legon, Jamu K. Alford, Hubert H. Lim

## Abstract

Ultrasound (US) can noninvasively activate intact brain circuits, making it a promising neuromodulation technique. However, little is known about the underlying mechanism. Here, we apply transcranial US and perform brain mapping studies in guinea pigs using extracellular electrophysiology. We find that US elicits extensive activation across cortical and subcortical brain regions. However, transection of the auditory nerves or removal of cochlear fluids eliminates the US-induced activity, revealing an indirect auditory mechanism for US neural activation. US likely vibrates the cerebrospinal fluid in the brain, which is continuous with the fluid in the cochlea via cochlear aqueducts; thus, US can activate the ascending auditory pathways and other non-auditory regions through cross-modal projections. This finding of a cochlear fluid-induced vibration mechanism challenges the idea that US can directly activate neurons in the intact brain, suggesting that future US stimulation studies will need to control for this effect to reach reliable conclusions.

## Introduction

Multiple neuromodulation techniques, such as deep brain stimulation (DBS), transcranial magnetic stimulation (TMS), and transcranial direct current stimulation (tDCS) have been used for treating various brain disorders, including tremors, depression, seizures, schizophrenia, pain, and tinnitus (Hallett, 2000; Johnson et al., 2013; Nitsche et al., 2008; Perlmutter and Mink, 2006). However, DBS and other invasive approaches have risks associated with surgery and involve high costs. TMS, tDCS and other noninvasive approaches do not require surgery, but they do not sufficiently achieve targeted activation (Brunoni et al., 2011; Deng et al., 2013; Markovitz et al., 2015a). More recently, ultrasound (US) stimulation has emerged as a potential approach that can address the trade-offs faced by modern neuromodulation technologies, in that it can be applied noninvasively but with the ability to activate or modulate targeted brain regions. For example, using indirect methods for measuring targeted neural activation, such as electroencephalography or perceptual tests, US has shown to activate or modulate primary somatosensory cortex (SC1) and primary visual cortex in humans or animals (Fry et al., 1958; Lee et al., 2015, 2016; Legon et al., 2014; Yoo et al., 2011). Primary motor cortex also appears to be activated with US stimulation in animals based on induced body or limb movements and electromyography measurements (King et al., 2014; Tufail et al., 2010; Ye et al., 2016; Yoo et al., 2011), though similar motor movements have not yet been achievable in humans.

There is growing interest and research in US neuromodulation, with an increasing number of publications in recent years. However, to our knowledge, there are still no in *vivo* mapping studies of neural responses within the brain that have confirmed that US is directly and locally activating neurons. In this study, we directly and simultaneously recorded across multiple locations within the brain, including SC1, primary auditory cortex (A1) and auditory midbrain using multi-site electrode arrays in an in vivo guinea pig preparation, and characterized the spatial neural activation pattern in response to US stimulation targeted at SC1 or A1.

Using similar US stimulation parameters and levels published in previous studies (Bystritsky et al., 2011; Mehić et al., 2014; Ye et al., 2016), we observed extensive activation across different brain regions, including A1 and SC1, with activation patterns that were unexpectedly consistent across different US stimulation locations, even when placing the transducer on the skull far from our recording location, or on the eyeball. More surprisingly, bilateral transection of the auditory nerves or removal of fluids from both cochleas caused the US-induced activity to disappear. These findings indicate that US stimulation across a wide range of parameters is activating the cochlea, which in turn activates the ascending auditory pathway up to A1 and possibly other cortical areas through cross-modal projections (Aitkin et al., 1981; Clemo et al., 2007; Foxe et al., 2000; Gruters and Groh, 2012; Murray and Wallace, 2011; Ramachandran and Altschuler, 2009; Schofield et al., 2011; Sigrist et al., 2013; Stein and Stanford, 2008). Cochlear vibration and activation may be achieved through US-induced vibration of the cerebrospinal fluid (CSF) in the head, which is continuous with the fluid in the cochlea via cochlear aqueducts (Gopen et al., 1997; Sohmer and Freeman, 2004; Sohmer et al., 2000). Combined with the results presented in the companion paper by Sato et al. (https://doi.org/10.1101/234211) investigating US-induced neural activity and motor responses in intact and deafened mice, these findings reveal an unexpected auditory activation effect caused by US stimulation, and the critical need for further studies to accurately characterize the direct neuromodulation capabilities of US.

## Results

### Indirect Activation of Central Auditory Circuits Using Pulsed Ultrasound

We first examined if pulsed US could directly activate the auditory cortex, since previous studies already reported that US could activate motor, visual, and somatosensory cortices (King et al., 2014; Lee et al., 2015, 2016; Tufail et al., 2010; Ye et al., 2016; Yoo et al., 2011). We recorded multi-unit spike activity (displayed as post-stimulus time histograms, PSTHs) and local field potentials from A1 with a 32-site electrode array while stimulating A1 with pulsed US (0.22 MHz, 100 kPa, 0.1 msec pulse duration (PD), single pulse, 500 msec trial duration (TD); Figure 1A). The transducer was attached to a plastic focusing cone filled with degassed deionized water, and the cone tip was coupled to the brain with degassed agarose. The US-evoked auditory responses were observed across all 32 sites (Figure 1C). The spike and negative-peak LFP responses to US occurred within 50 msec of the stimulus onset, which is consistent with the time scale of neural activity in primary motor cortex from a previously published study (Tufail et al., 2010). US-evoked auditory responses could be achieved with numerous US parameters (Table S1).

**Figure 1.**
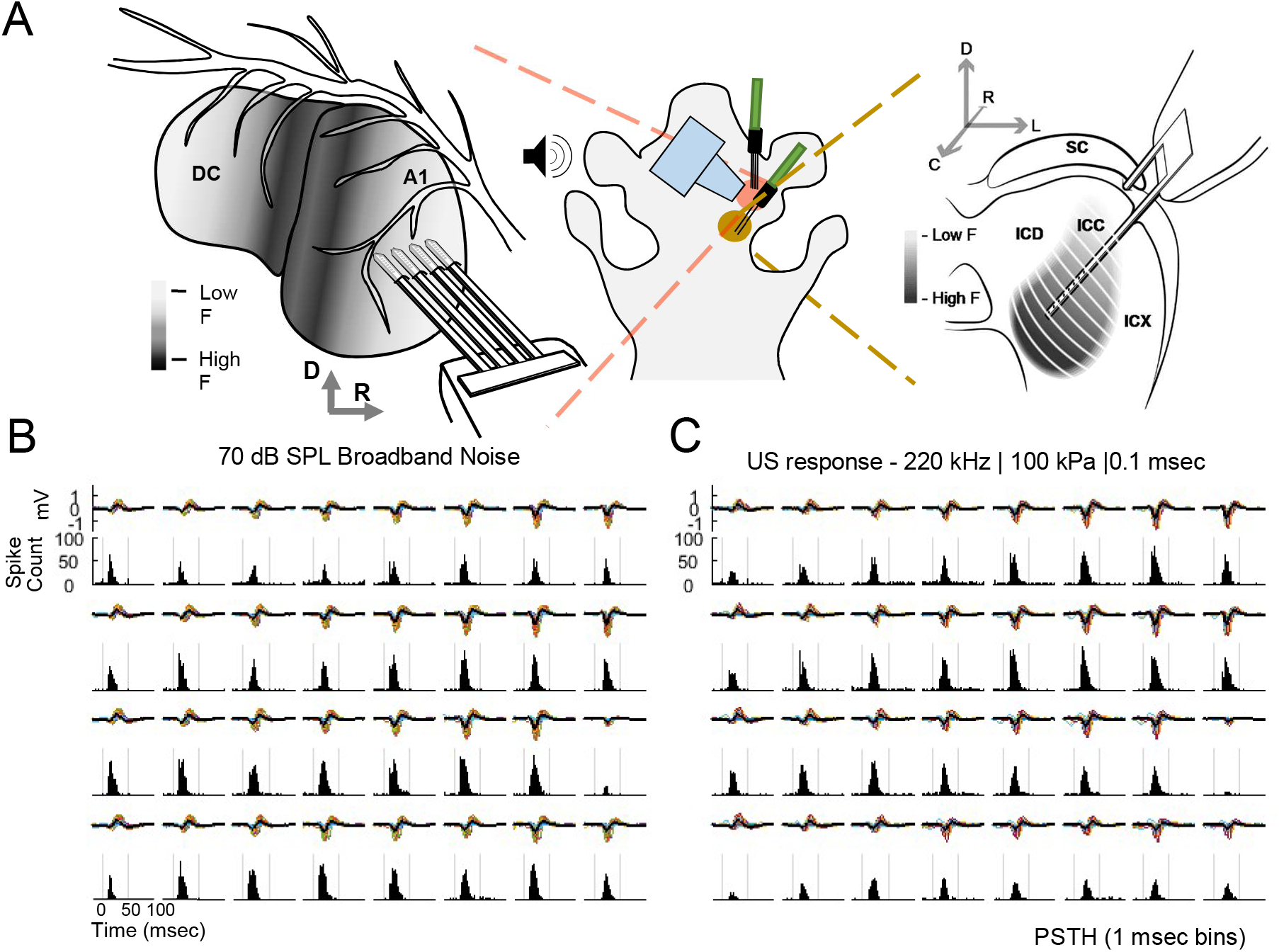
A1 responses to US stimulation and broadband noise (BN). **A** Two 32-site electrode arrays were inserted into the right ICC and A1 of anesthetized guinea pigs, and both probes were positioned such that recorded neurons were tuned to frequencies that spanned a wide range. The US transducer was placed over the exposed A1 (coupled with agar), and acoustic stimulation was presented to the left ear through a hollow ear bar. **B** A1 responses to 70 dB SPL BN acoustic stimulation are shown. For each recording site (i.e., subplot), the local field potentials (LFPs) across 100 trials are plotted in the top portion and the post-stimulus time histograms (PSTHs) of spiking activity are plotted in the bottom portion. Each row corresponds to the linearly spaced electrode sites along one shank of the 4-shank probe. Activity is observed on all 32 sites. Time is relative to stimulus onset. **C** A1 responses to US stimulation (220 kHz, 100 kPa, 0.1 msec PD, 500 msec TD) are shown. Similar to BN responses in **B**, activity is observed on all 32 sites.

Surprisingly, the US-induced responses across sites closely resembled the activity evoked by audible broadband noise at 70 decibels sound pressure level (dB SPL; Figure 1B). We questioned if the animal was sensitive to ultrasound being directly generated by the transducer or was hearing audible sound during US application, even though we could not hear any sound originating from the US transducer. As shown in Figure 2B, we placed the transducer on US gel that was unconnected to the animal’s body and recorded from A1 while presenting US stimulation (220 kHz, 100 kPa, 10 msec PD, 500 msec TD). We could not elicit any US-induced activity in A1, even when placing the transducer closer to the animal’s ears. In contrast, A1 activity was clearly observed whenever the transducer was coupled to the animal’s head or brain (Figure 2A; also see Figure 1C). Unexpectedly, the transducer could be positioned anywhere on the head or body, even on the eyeball, and still elicit strong and similar patterns of A1 activity as long as the transducer was directly coupled to the animal (Figure 2D–H). These results contradict the localized brain activation pattern that would be expected from the focused energy profile achievable with US (Figure S1).

**Figure 2.**
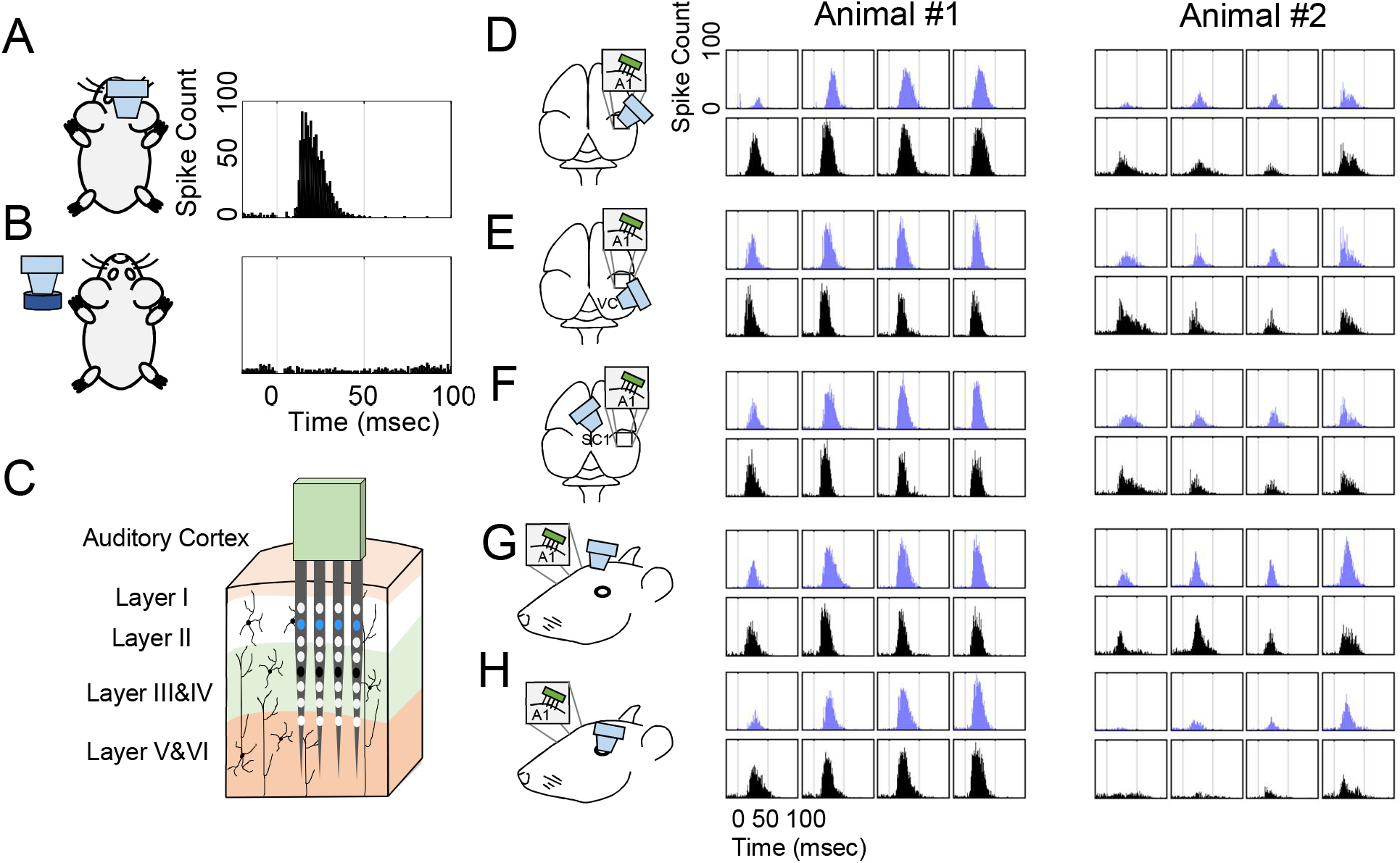
A1 responses to US stimulation of different regions in the brain, on the head, and coupling gel detached from the body. **A** US stimulation (220 kHz, 100 kPa, 10 msec PD, 500 msec TD) of A1 elicited spike activity over all 32 sites. Only one typical site is shown here. **B** US stimulation of coupling gel detached from the animal’s body with same US parameters in **A** does not elicit any A1 spike activity over all 32 sites, confirming the responses in **A** are not attributed to an external sound generated by the US transducer. The same site in **A** is shown here. **C** Diagram of Layer II (blue) and Layer III/IV (black) of auditory cortex is shown, which corresponds to the blue and black PSTHs shown in **D-H. D-H** Diagrams and PSTHs representing spike responses to US stimulation (220 kHz, 100 kPa, 10 msec PD, 500 msec TD) of exposed auditory cortex (**D**), exposed visual cortex (**E**), exposed somatosensory cortex (**F**), the left side of the intact skull (**G**), and the left eyeball (**H**) in two animals are shown. All stimulation locations result in A1 activation of 64 sites across the two animals and are surprisingly similar in each animal even though the stimulation targets were quite different. All US setups in **D-H** were coupled to the animal with agar. Time is relative to stimulus onset.

In two animals we also recorded simultaneously from A1 and from the central nucleus of the inferior colliculus (ICC; Figure 1A), which is the main ascending region of the auditory midbrain that then projects up to the thalamus and A1. The original motivation for recording from ICC in response to US stimulation of A1 was to characterize the spatial spread of US activation in A1. A previous study from our lab showed that electrical stimulation of A1 activates descending cortical-to-midbrain feedback pathways to ICC that are spatially organized in a topographic pattern, in which A1 neurons sensitive to a given pure tone frequency activate ICC neurons that are sensitive to a similar pure tone frequency (Markovitz et al., 2013). Thus, it is possible to record across this tonotopic gradient of the ICC while stimulating a specific region of A1 and use the ICC read-out to assess the spread of activation across A1. Stimulation with US of different A1 locations elicited extensive activity throughout ICC with no indication of localized effects. Unexpectedly, the first spike latencies in ICC were much shorter than those in A1, which contradicts what would be expected if we were directly activating A1 that then activated descending projections to ICC (Figure 3). This unexpected result occurred regardless of the US stimulation parameters, in which the mean first spike latencies across US parameters were significantly shorter for ICC compared to A1 (5.43 ± 0.15 versus 17.96 ± 1.35 msec, *P* ≪ 0.001). Furthermore, the first spike latencies observed for A1 and ICC in response to US stimulation approximately resembled those for audible broadband noise stimulation (Figure 3, rightmost bars), suggesting US may be activating the ascending auditory pathway at the peripheral or cochlear level.

**Figure 3.**
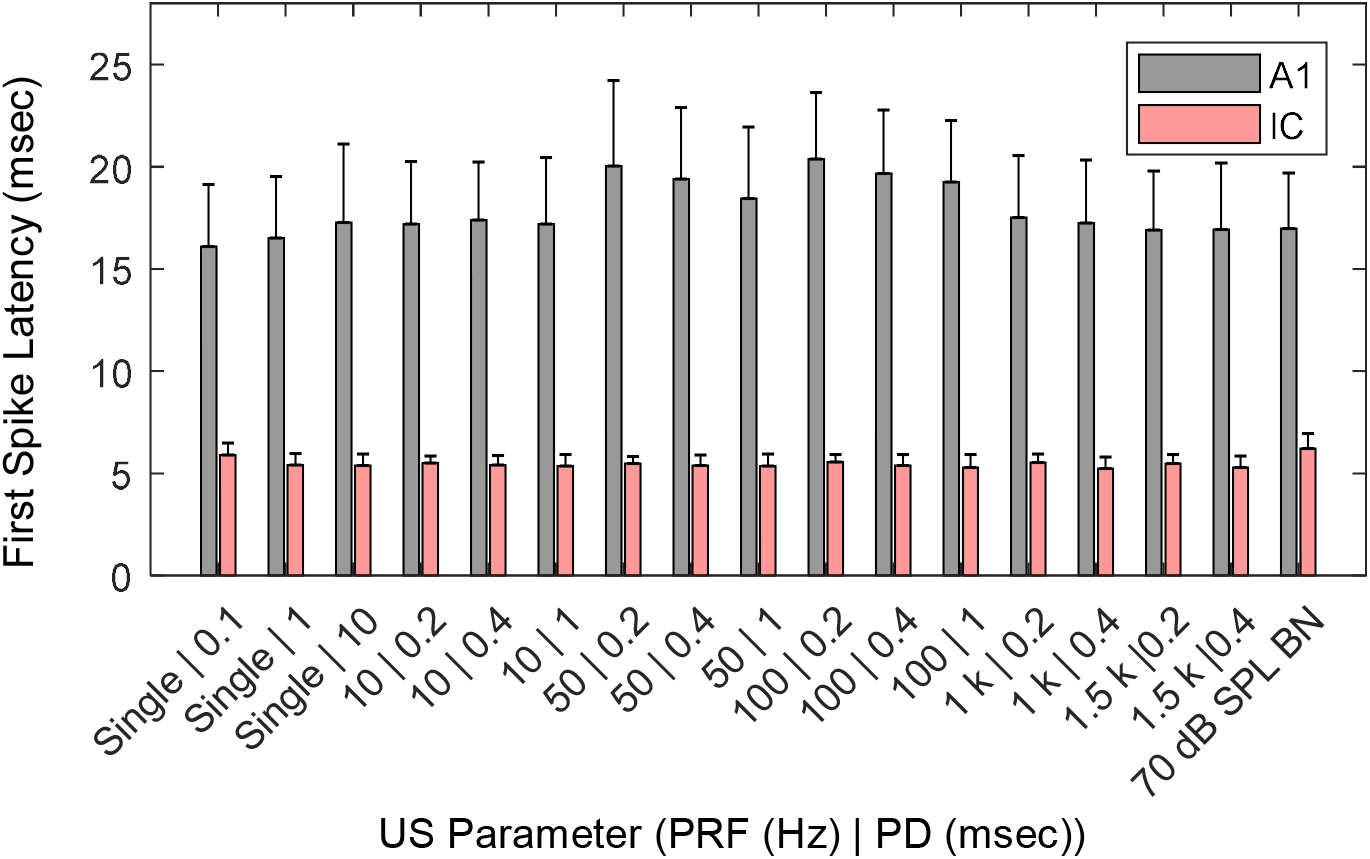
First Spike Latencies of A1 and ICC in response to US stimulation of A1. First spike latencies of ICC and A1 responses to US stimulation of exposed A1 at 200 kPa are shown. One condition of 70 dB SPL BN acoustic stimulation is also shown at the far right of the plot for comparison. For each stimulation parameter, latencies were averaged across all recording sites and plotted with standard error bars. In general, IC neurons had shorter latencies than A1, suggesting that ICC was activated before A1, similar to what would be expected for acoustic-driven activation along the ascending auditory pathway. US-induced latencies were similar to those of BN stimulation, indicating that US may be activating the auditory system early in the ascending auditory pathway. Data are represented as mean ± SD, which is pooled from 91 recording sites across 2 animals.

To confirm that the US-induced activity is caused by peripheral or cochlear activation, we performed control experiments in which we characterized US-induced activity in A1 before and after transection of both auditory nerves. Any auditory activity originating from the cochlea to the brain (i.e., within the brainstem, midbrain, thalamus and cortex) must pass through the auditory nerves. Figure 4A and 4B show A1 activity elicited from broadband noise and US stimulation, respectively, in which similar and strong responses across nearly all sites are observed for both stimulation conditions before transection. Note that an US pressure level of 100 kPa for a 10 msec PD and 500 msec TD elicits a strong response similar to what is observed for a 70 dB SPL broadband noise stimulus, which is considered a loud sound in guinea pigs based on their thresholds of hearing. After transection of the auditory nerves, all of the stimulus-driven activity disappeared for both stimulation conditions (Figure 4C, D). We did not observe any A1 activity in response to US stimulation after transection of the auditory nerves for a wide range of parameters, except in one animal. Figure 4F is an example in one animal using a high pressure of 2 MPa where we observed a statistically significant response on one site (red box) after nerve transection. Other US parameters at high pressures in that same animal were able to elicit activity on one or a few sites (Figure 4E and Table S2). Considering the inability to repeat these results in any other animal and the long onset latencies of the US-induced responses after nerve transection (i.e., still between 11–16 msec), we believe the auditory nerves may not have been fully transected in that one animal. Overall, the results from these control experiments confirm that all or nearly all of the activity observed in A1 in response to US is caused by a peripheral auditory pathway passing through the auditory nerve to the brain. Later, we show that this peripheral pathway requires the fluids within the cochlea. As to whether direct brain activation is possible using different US parameters than what was used in our study requires further investigation.

**Figure 4.**
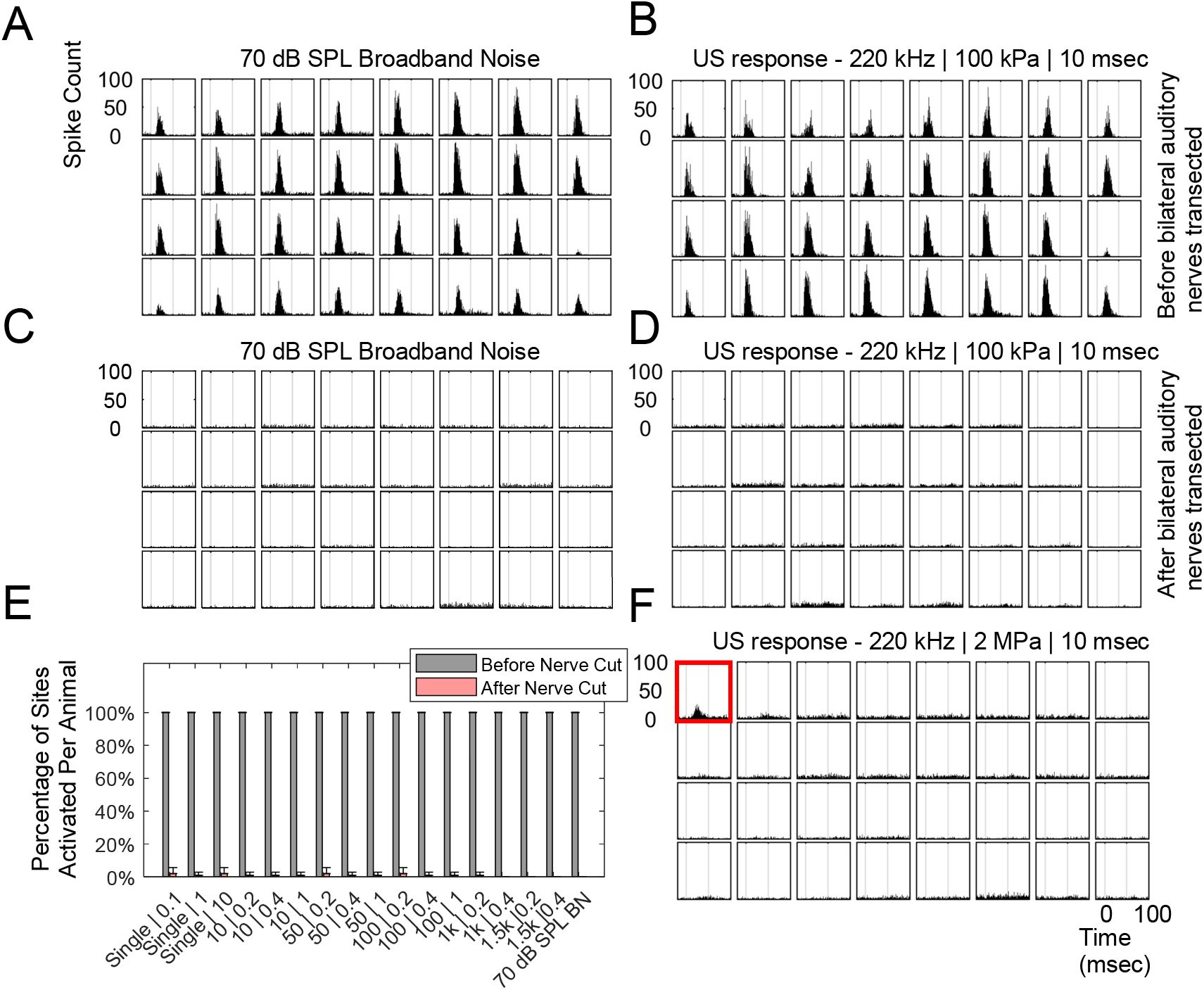
US-evoked A1 responses are eliminated with bilateral transection of auditory nerves. **A-B** A1 spike responses to BN stimulation (**A**; 70 dB SPL) and US stimulation of exposed A1(**B**; 220 kHz, 100 kPa, 10 msec PD, 500 msec TD) before transecting the auditory nerves on both sides. Nearly all 32 sites of the recording probe show strong spiking activity. **C-D** A1 recordings in response to BN and US stimulation were repeated after bilateral transection of the auditory nerves as shown in **C** and **D** respectively. In both cases, there is no apparent stimulus-driven activity. **E-F** When increasing US pressure up to 2 MPa, there were only less than 5% of sites activated per animal (n=3; total number of sites per condition was 95) that responded to US stimulation of A1 after bilateral transection of the auditory nerves. Only one animal out of three showed residual activity after nerves were cut, and only for a small number of sites (example shown in **F**). See also Table S2. Time is relative to stimulus onset. Data are represented as mean ± SD.

### Indirect Activation of Somatosensory Cortex Using Pulsed Ultrasound

Since previous studies demonstrating US-induced activity in the brain targeted other cortical regions than the auditory cortex, we questioned whether our findings concerning auditory activation were specific to A1. We performed several experiments targeting somatosensory cortex with US, since multiple studies have shown or suggested that US can evoke or modulate spike activity in SC1 and supplementary eye field (Lee et al., 2015; Legon et al., 2014; Wattiez et al., 2017). We implanted a 4-shank, 32-site electrode array into SC1 and recorded multi-unit spike activity in response to pulsed US targeted towards SC1 (200 kPa, 1 kHz pulse repetition frequency (PRF), 0.5 msec PD, 20 pulses, 6 sec TD; Figure 5A). As shown in Figure 5E, US-evoked SC1 activity was observed on 31 out of 32 sites (based on statistical analysis described in the Methods) in one animal. The raster plot for a typical site (#29 in Figure 5E) shows that the spikes were consistently observed with a wide temporal range across 100 trials. Results across multiple animals show that US could reliably elicit spike activity in SC1 with a mean percentage of activated sites of 60.3 ± 23.8% across seven animals (Figure 5B). A wide range of US parameters we tested could elicit activity in SC1 (Table S3).

**Figure 5.**
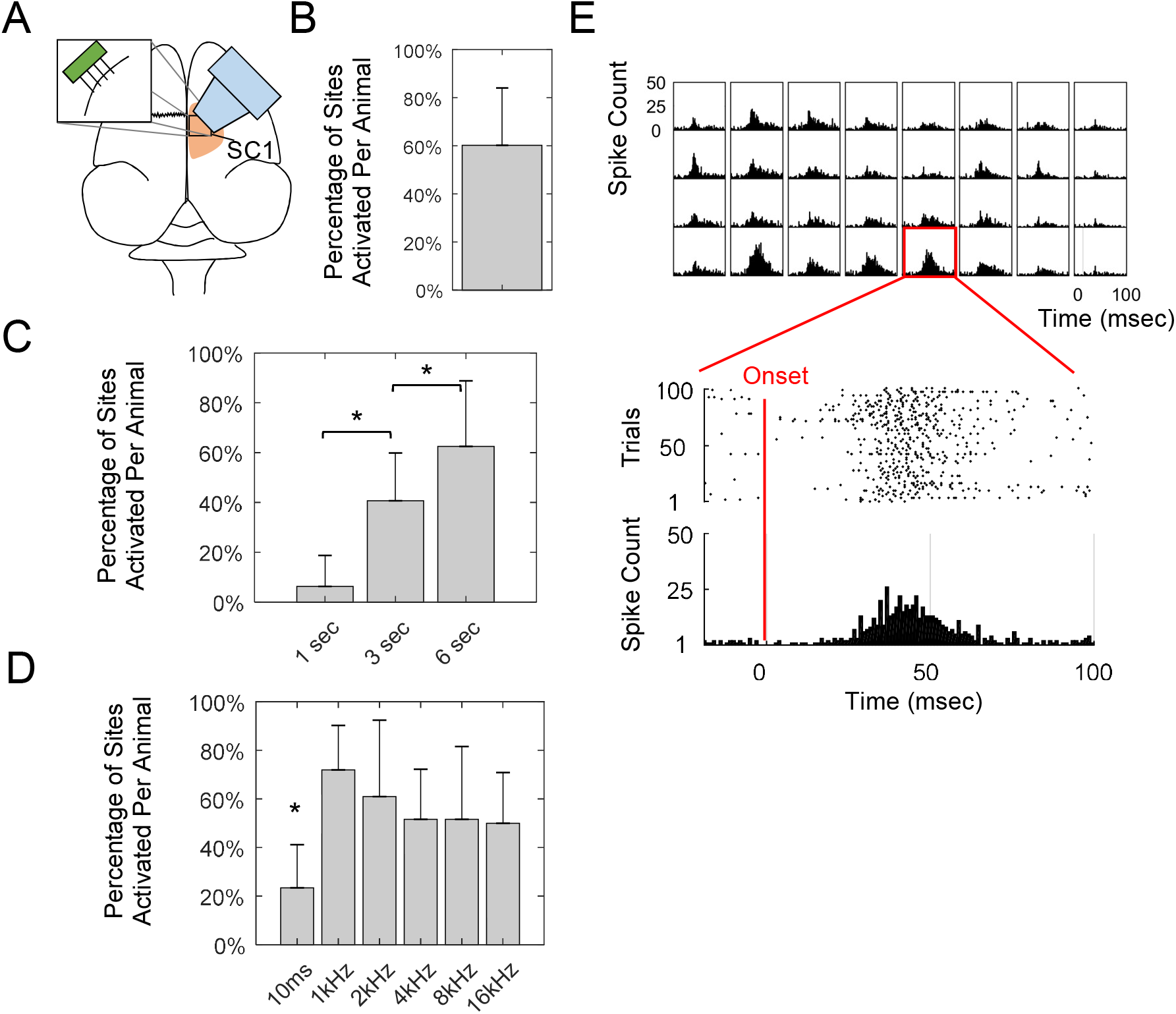
US stimulation of SC1 with different US paradigms. **A** 32-site electrode arrays were inserted into the right SC1 of anesthetized guinea pigs. The US transducer was placed over SC1 coupled to the brain with agar. **B** On average, 60.3% of SC1 recording sites were activated per animal (n=7) with US stimulation (200 kPa, 1 kHz PRF, 0.5 msec PD, 20 pulses, 6 sec TD). **C** The percentage of SC1 sites activated per animal (n=4) increased (two-tailed, unequal variance, ranked t-test; \**P* < 0.05) as trial duration increased, reaching 62.5% of sites with a 6 sec TD in this cohort of animals. **D** PRF also affected the percentage of sites activated, in which repeated US pulse paradigms with different PRFs activated more sites than a single 10 msec pulse US stimulus (two-tailed, unequal variance, ranked t-test; \**P* < 0.05). Lower PRFs appeared to activate more sites. **E** A typical example of SC1 activation across a 32-site electrode array in response to US stimulation (200 kPa, 1 kHz PRF, 0.5 msec PD, 20 pulses, 6 sec TD). One site example is magnified for better visibility, showing a PSTH and a raster spike plot (i.e., dots corresponding the time occurrence of each spike for each trial of US stimulation). Time is relative to stimulus onset. Detailed US paradigms used in **C** and **D** can be found in Table S3. Data are represented as mean ± SD.

To investigate US-induced activation effects similar to what was explored in previous studies (Lee et al., 2015, 2016; Tufail et al., 2010), we specifically tested US paradigms with different TDs of 1, 3, and 6 sec and PRFs of 1, 2, 4, 8, and 16 kHz (in addition to a single pulse), while keeping other parameters unchanged. With a shorter TD of 1 sec, US only elicited 12.5 ± 6.3% of activated sites, which was significantly lower (*P* < 0.05) than 40.6 ± 19.3% for 3 sec TD and 62.5 ± 26.3% for 6 sec TD across four animals (Figure 5C). This finding is consistent with previous US studies stimulating the somatosensory, motor and visual cortices in which longer TDs caused greater activation (Lee et al., 2015, 2016; Tufail et al., 2010). For US stimulation with different PRFs (1, 2, 4, 8 and 16 kHz), single pulse stimulation (10 msec PD, 1 pulse, 6 sec TD) elicited a significantly lower percentage of activated sites across four animals (*P* < 0.05; 23.4 ± 17.8% versus 71.9 ± 18.4%, 60.9 ± 31.5%, 51.6 ± 20.7%, 51.57 ± 29.7%, and 50.0 ± 20.9%, respectively; Figure 5D), indicating that single pulse stimulation is not as effective in eliciting SC1 activity as repeated pulse patterns with varying PRFs. Although not significant (P > 0.05), there appears to be a trend of greater activation with lower PRFs down to 1 kHz, which may explain why previous US studies stimulated the cortex using low PRFs such as 0.5 and 1.5 kHz, rather than trying single pulse US (Lee et al., 2015, 2016; Tufail et al., 2010).

Since we have shown that US-evoked A1 activity was elicited through an ascending auditory pathway originating at the peripheral or cochlear level and that US stimulation of different head or brain locations elicited similar responses in A1, we also questioned whether similar unexpected results would be observed for SC1. In the next section, we present data on the effects of eliminating cochlear function on US-induced SC1 activity. Here, we investigated the effects of US stimulation of a different cortical region, particularly visual cortex, on the neural response patterns elicited in SC1 (Figure 6A). Similar to what we observed for US-induced activity in A1, we were still able to elicit strong SC1 activation while stimulating a distant location with US, even though the focal zone in visual cortex was approximately 10 mm away from the SC1 recording sites. Thus, it is unlikely that US stimulation was directly activating SC1 since the target zone was far from SC1 (see Figure S1 for US energy field profile). The onset latency of neural activity was generally greater than 20 msec (Figure 6D,E), which is longer than would be expected for direct activation of cortical neurons with US. Surprisingly, the onset latency of neural activation when stimulating directly over SC1 versus visual cortex was not noticeably different (Figure 5E versus Figure 6D,E) and these longer ranges of latencies are consistent with those published in previous studies for US-induced activation of motor cortex (Tufail et al., 2010, 2011).

**Figure 6.**
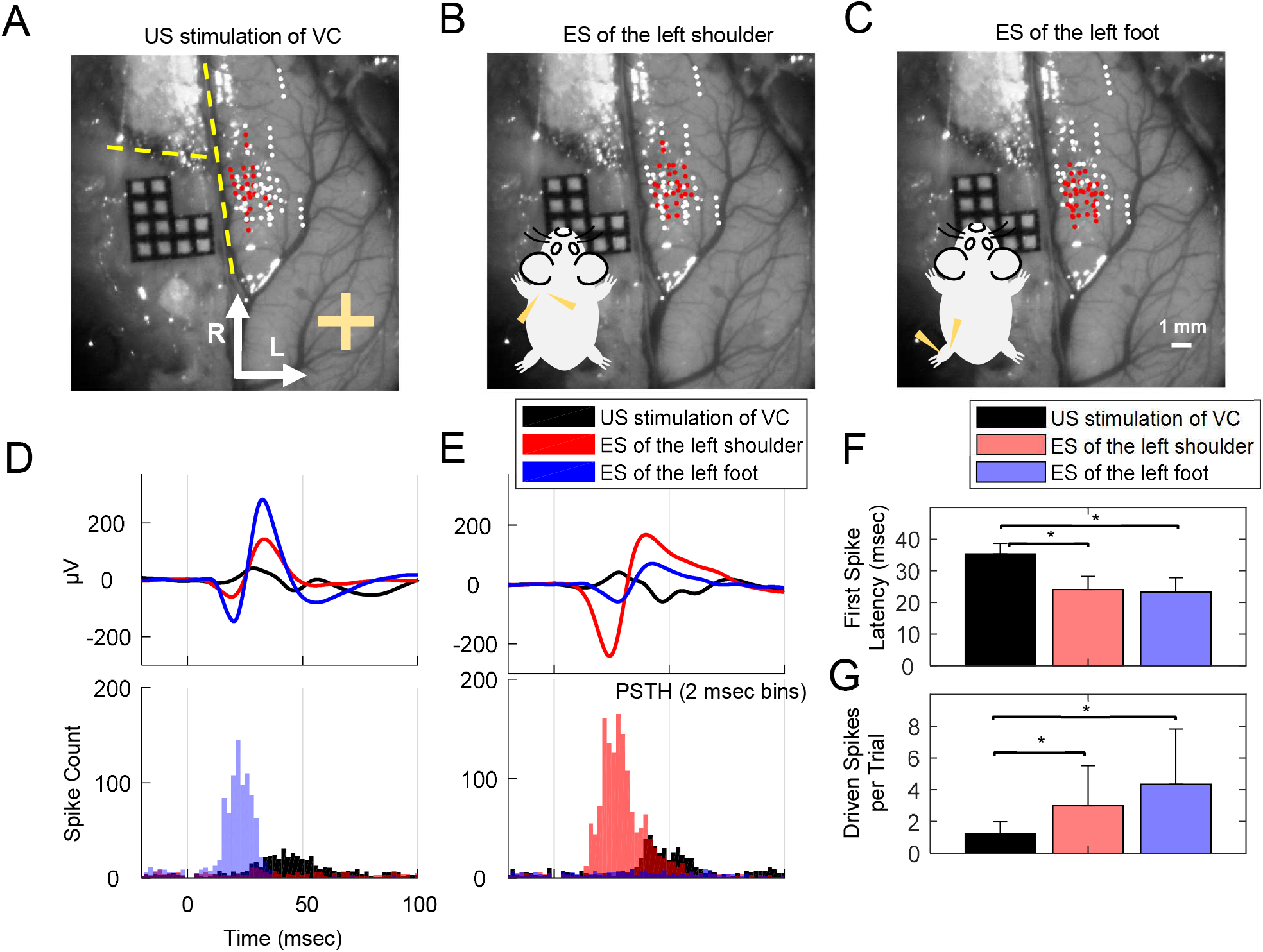
Activation of SC1 with US stimulation of visual cortex. **A-C** Locations of recording electrodes in SC1 that exhibited activity (red dots; white dots exhibited no activity) in response to US stimulation (400 kPa, 1 kHz PRF, 2.5 msec PD, 20 pulses, 6 sec TD) of the exposed visual cortex (**A**; plus sign is location of visual cortex, VC), electrical stimulation (ES) of the left shoulder (LS; **B**), and ES of the left foot (LF; **C**). Yellow dotted lines indicate the locations of the midline and the Bregma suture line. Red dots represent locations where at least 50% of active recording sites along an inserted electrode shank were activated. ES was presented in bipolar mode with two subcutaneous needle electrodes with a single, biphasic, cathodic-leading pulse (205 μs/phase, levels of 2.82 mA). US transducer was coupled to VC with agar. Data is pooled across 4 animals and 576 recording sites, where 225 (39.1%), 221 (38.4%), and 300 (52.1%) sites were activated by US stimulation of VC, ES of the LS, and ES of the LF, respectively. Out of the 225 sites responding to US stimulation of VC, 105 (46.7%) and 136 (60.4%) sites overlapped with the sites responding to ES of the LS and ES of the LF, respectively. **D-E** Examples of local field potentials (LFPs) and PSTHs in response to US stimulation of VC (black), ES of the LS (red), and ES of the LF blue). For **D**, ES of the LF elicited the most spikes and largest LFPs, while ES of the LS elicited the most spikes and largest LFPs in **E. F** On average, first spike latencies were much longer (two-tailed, unequal variance, ranked t-test; \**P* < 0.05) in SC1 for US stimulation of VC (35.29 ± 3.39 msec) than for ES of the LS (24.07 ± 4.22 msec) and ES of the LF (23.31 ± 4.48 msec). **G** On average, driven spike counts were also lower (two-tailed, unequal variance, ranked t-test; \**P* < 0.05) for US stimulation of VC (1.23 ± 0.76 spikes per trial) compared to ES of the LS (2.99 ± 2.53 spikes per trial) and ES of the LF (4.35 ± 3.48 spikes per trial). Data are represented as mean ± SD.

To better interpret the extent and delay of activation caused in SC1 in response to US of visual cortex, we compared the activity to what is elicited when electrically stimulating a somatosensory pathway, such as through electrical stimulation of a body region. We electrically stimulated the left shoulder or left foot and recorded neural activity across multiple locations in the contralateral (right) SC1 (Figure 6B,C). US stimulation targeted at the visual cortex still caused activation across a spatially broad region of SC1 (Figure 6A), but tended to elicit weaker and longer latency responses than direct electrical stimulation of the somatosensory pathway (Figure 6D–G). Combined with the activation latencies observed in Figure 5, these data indicate that US was unlikely causing direct activation of the cortex and the neural activation mechanism was mostly insensitive to location of stimulation on the head, which is consistent with the findings shown in Figure 2.

### Removing a Cochlear Fluid Pathway Eliminates Auditory and Somatosensory Activity Induced by Ultrasound Brain Stimulation

Since the US-induced activity in A1 or SC1 appeared to be insensitive to location of stimulation across the head and was not altered substantially if US was applied to the skull or to soft tissue such as the brain surface or eyeball, we hypothesized that the mechanism of activation occurs through a fluid pathway by vibrating the CSF in the head (Figure 7A). The CSF in the brain is continuous with the fluid in the cochlea through the cochlear aqueducts (i.e., fluid channels between the head and cochlea; Gopen et al., 1997; Sohmer and Freeman, 2004; Sohmer et al., 2000). Therefore, vibrations of the CSF in the head or brain, such as through US stimulation, could induce vibrations in the cochlear fluids. Cochlear vibrations would then activate the hair cells along the cochlea that then innervate the auditory nerve fibers to the ascending auditory pathway in the brain (Moore et al., 2010; Yost, 2000). Activation of different auditory nuclei can lead to activation of non-auditory regions, such as SC1, through cross-modal projections consistent with the range of delays observed in Figures 5 and 6.

**Figure 7.**
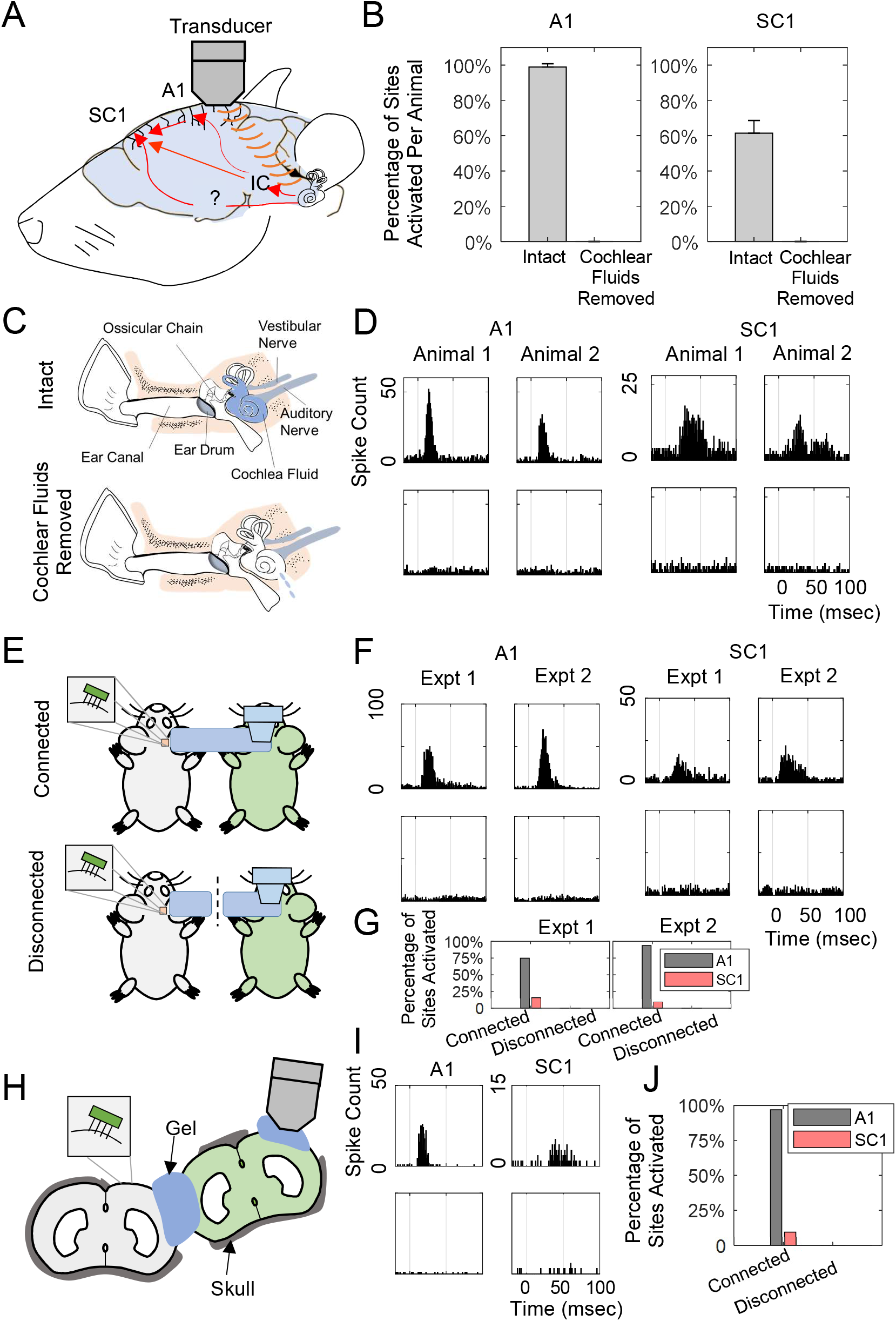
Removing bilateral cochlear fluids eliminates US-induced auditory or somatosensory activity. **A** Illustration of the cochlear fluid pathway. US may be vibrating cerebrospinal fluids (CSF) that then vibrates cochlear fluids (via cochlear aqueducts; i.e., CSF and cochlear fluids are part of a continuous fluid medium) to induce activity along the ascending auditory pathway, which also has multimodal projections to somatosensory nuclei. **B** The percentages of A1 and SC1 sites activated by US stimulation before removal of cochlear fluids per animal were 99.0 ± 1.8% and 61.5% ± 7.2%, respectively. In both cases, all activity was eliminated after fluids were removed in all animals (n= 3). **C-D** The illustration shows the peripheral auditory system before (upper plots in **C**) and after removal of the cochlear fluids (lower plots in **C**). In two different experiments, A1 activity elicited by US stimulation of the exposed A1 (50 kPa, 10 msec PD, 500 msec TD) is shown in **D** (upper PSTHs), in which all activity was eliminated after cochlear fluids were removed (lower PSTHs). Similar findings are shown for SC1 activity in response to US stimulation of exposed SC1 (200 kPa, 1 kHz PRF, 0.5 msec PD, 20 pulses, 6 sec TD). **E-F** The illustration shows how US can indirectly activate the A1 (100 kPa, 10 msec PD, 500 msec TD) and SC1 (400 kPa, 1 kHz PRF, 0.5 msec PD, 20 pulses, 6 sec TD) of one animal (grey) by transcranially stimulating the brain of another animal (green). A fluid channel made of US gel connects the exposed cortex of grey animals and the skull of the green animal (upper) to enable this animal-to-animal activation, as shown in the upper plots of **F**. Breaking this gel coupling (dotted line), eliminates the animal-to-animal activation (lower plots in **F**), further confirming a fluid/gel mechanism of activation of the brain with US stimulation. Data from two experiments are shown. G Bar plots illustrate that A1 and SC1 sites activated in grey animal (upper plots in **E**) were 75.0% and 15.6% in experiment #1, and 93.7% and 10.0% in experiment #2, which were eliminated after breaking of the gel channel connecting two animals (lower plots in **E**). **H-I** The illustration shows how the gel connects the exposed brain of the live animal (grey) and the left exposed brain of the dead animal (green). US stimulation of the right exposed brain of the green animal activates the A1 (200 kPa, 10 msec PD, 500 msec TD) and SC1 (400 kPa, 1 kHz PRF, 0.5 msec PD, 20 pulses, 6 sec TD) of the grey animal (upper plots in **I**), which can be eliminated by breaking the gel channel connecting the two brains (lower plots in **I**). **J** Bar plots illustrate that A1 and SC1 sites activated in grey animal were 96.9% and 9.38% when connected with green animal through gel, which reduced to 0% after breaking the gel channel. Data are represented as mean ± SD.

To confirm our hypothesis, we developed a method for removing the fluids from the cochlea using a syringe inserted through the round window membrane (Figure 7C), as further described in the Methods. Figure 7B,D (also Figure S2) shows that extensive US-induced activity is possible in A1 and SC1, but disappears after removal of the cochlear fluids, confirming our hypothesis that the mechanism of US activation requires a cochlear fluid pathway. Inspired by a previous fluid conductive hearing study connecting the CSF of two animals through a water tube (Sohmer and Freeman, 2004), we designed a similar two-animal setup (Figure 7E) in which US was applied to a dead animal (green) and neural activity was recorded from A1 or SC1 of a live animal (grey). The right skull of the live animal was opened to expose the brain and covered with an agar layer. A fluid channel made of US gel connected the brain of the live animal to the skull of the dead animal. US was applied to the right skull of the dead animal. US stimulation of the dead animal elicited activity in A1 and SC1. Consistent with a fluid mechanism, the US-induced activity disappeared once the gel channel was broken (Figure 7F,G). To confirm that a fluid mechanism occurred within the brain, and not only through the gel channel from the dead animal, we performed another control experiment in which we directly connected the exposed brains of both animals, as shown in Figure 7H, and stimulated a different part of the exposed brain in the dead animal that was not in contact with the gel channel, thereby requiring conduction through the brain. Figure 7I,J shows that US can vibrate or travel through the brain of the dead animal to elicit neural activation in the live animal via a gel channel, and this activity disappears once the gel channel is broken. We used a dead animal for US stimulation to eliminate any conductive activation effects through the gel channel that may occur from US activity elicited in a live animal. Overall, these control experiments shown in Figure 7 confirm a fluid mechanism through the brain and cochlea for US-induced activation of the brain.

## Discussion

Our study revealed that US is able to activate A1 with spatial-peak pulse-average acoustic intensities (*I*_SPPA_) as low as 20 mW/cm^2^ (~25 kPa for the 0.22 MHz transducer), which are lower than what has been previously published for brain activation (Bystritsky and Korb, 2015; Bystritsky et al., 2011; Tufail et al., 2010). However, this US-evoked A1 activity is through a non-direct cochlear fluid pathway rather than direct activation of A1 neurons. This finding is supported by multiple pieces of evidence. First, US stimulation of other non-auditory regions far from A1 (e.g., exposed SC1, contralateral skull or eyeball) caused similar activation of A1 as that from directly targeting A1. Second, US stimulation of A1 elicited strong activity in ICC, but the onset latencies of activity in ICC were significantly shorter than those recorded in A1 (*P* ≪ 0.001), suggesting ICC activation was not elicited through the descending corticofugal projections from A1. In addition, the average onset latency of US-induced activity in A1 was approximately 14 msec, which is similar to the latency required for acoustic stimuli to vibrate the cochlea and activate the auditory nerve up to the brainstem, ICC, thalamus, and A1 through the ascending auditory pathway in guinea pigs (14.4 ± 4.4 msec; Wallace et al., 2000). Finally, the US-evoked A1 activity could be eliminated by either transecting the auditory nerves or removing the cochlear fluids for both ears.

We confirmed that US can activate SC1 with a wide range of parameters with intensities as low as 80 mW/cm^2^ *I*_SPPA_ (~50 kPa for the 0.22 MHz transducer), which is on the same order of magnitude as previous studies (Lee et al., 2015; Mehić et al., 2014; Tufail et al., 2010; Ye et al., 2016; Yoo et al., 2011). US stimulation of non-SC1 regions such as the visual cortex was also able to activate SC1. Similar to the findings for US-evoked A1 activity, we found that US-evoked SC1 activity was eliminated by removing cochlear fluids in both ears, suggesting US does not directly activate SC1 neurons but instead causes SC1 activation through a vibratory cochlear pathway that likely activates non-auditory neurons via cross-modal projections.

Numerous studies have shown extensive connections and interactions among different brain circuits (e.g., auditory, somatosensory, motor, visual, and high-level cognitive nuclei (Gruters and Groh, 2012; Ramachandran and Altschuler, 2009; Schofield et al., 2011; Sigrist et al., 2013; Wang et al., 2008)) to support this US-induced cross-modal mechanism. As to how much this US-induced cochlear mechanism contributed to the activation effects observed in previous US stimulation studies needs further investigation (e.g., Mehić et al., 2014; Tufail et al., 2010, 2011; Ye et al., 2016; Yoo et al., 2011).

One unexpected finding from our study is the large extent of activation across the auditory system that is possible with US stimulation at low intensities. For example, strong activation of A1 is possible at a pressure of 100 kPa (330 mW/cm^2^ *I*_SPPA_) for a 0.1 msec PD and 500 msec TD (e.g., similar activity to a 70 dB SPL broadband noise). Previous US studies have used intensities on the order of 1 W/cm^2^ *I*_SPPA_ for their neural activation or motor movement experiments (Mehić et al., 2014; Tufail et al., 2010; Ye et al., 2016; Yoo et al., 2011), which would be expected to elicit a very loud sound in the animals. In lightly anesthetized animals, it is possible that such loud US-induced sounds could elicit an auditory startle response and movements of limbs, whiskers or the tail (Geyer and Swerdlow, 2001; Grimsley et al., 2015; Turner et al., 2006). Previous studies have shown that US can induce motor movements, but usually in lightly anesthetized animals (Mehić et al., 2014; Tufail et al., 2010, 2011; Ye et al., 2016). Our results combined with those presented in the companion paper by Sato et al. (https://doi.org/10.1101/234211) raise the question as to whether the US-induced movements demonstrated in previous studies were caused by loud sounds through a vibratory cochlear effect induced from US stimulation. It is interesting that some studies found that US positioned on caudal locations of the head, closer to where the cochleas are located, achieved stronger and more reliable motor movements than US positioned over the motor cortices (Mehić et al., 2014; Ye et al., 2016). It is also interesting that the success rate for eliciting motor responses is quite low for different locations (usually less than 50%). Animals can experience variable responses and adaptation to the startle response in which it is more difficult to elicit startle reflexes over repeated or regular presentations (Davis, 1970; Glowa and Hansen, 1994; Grimsley et al., 2015), which may contribute to the high failure percentages of motor movements reported in previous US stimulation studies.

Although we find that removal of bilateral cochlear fluids eliminates US-evoked A1 and SC1 activity in guinea pigs, it should not be assumed that the same is true for other animal species. At least based on the consistent findings in mice in the companion paper by Sato et al. (https://doi.org/10.1101/234211), it appears that the US-induced activation of the brain via a cochlear pathway is valid for rodents. However, further studies are needed to assess the confounding effects of US-induced cochlear activation in other species with larger head sizes, including humans. Previous studies in humans have shown that US stimulation applied to the head with low frequencies (27–33 kHz) can elicit sound perception via a skull vibration mechanism (Ito and Nakagawa, 2010; Nakagawa et al., 2006), but the auditory activation effects for higher US frequencies used in neuromodulation studies still need to be investigated. There may also exist other US parameters that can achieve direct brain activation as well as modulation of neurons that were not tested in our study. There is evidence that US can directly activate neurons in in *vitro* experiments using brain slice cultures and modulate ion channels in oocytes (Kubanek et al., 2016; Tyler et al., 2008), which cannot be explained by a vibratory cochlear mechanism. There is also a recent study in humans demonstrating that US stimulation of the somatosensory cortex can elicit tactile sensations in the hand (Lee et al., 2015), which cannot be explained by a vibratory cochlear mechanism. Therefore, further studies are still needed to fully explore the parameter space to characterize the type and extent of US activation and modulation that is possible within the brain across species. Our findings reveal the critical need in these future studies to account for or eliminate confounding effects that can lead to artificial or indirect neural activation when applying US stimulation to the head. US-induced cochlear vibration is one confounding effect. Other confounding effects may include US-induced activation of skin receptors on the head or vibration of the skull or eyeballs, which in turn could activate various sensory and motor pathways that contribute to the overall activity of the brain.

## Acknowledgements

We would like to thank Daniel Zachs and Pooja Mehta for their help with the experimental setup and data interpretation. We also would like to thank Alyona Haritonova for contributions with ultrasound setup and technical discussions. This research was supported by SONIC Lab Discretionary Funds and a MnDRIVE Brain Conditions Innovations Grant led by Hubert Lim, Jamu Alford and Wynn Legon.

## Author Contributions

Conceptualization, H.H.L, H.G., M.H., J.K.A., S.J.O., and W.L.; Methodology, H.G., H.H.L., M.H., S.J.O., C.D.G., J.K.A., Y.K., and T.L.; Software and Formal Analysis, H.G., M.H., S.J.O., and H.H.L.; Investigation, H.G., M.H., S.J.O., T.L., C.D.G., and Y.K.; Writing – Original Draft, H.G., H.H.L., C.D.G., and M.H.; Writing – Review & Editing, H.G., H.H.L., S.J.O., C.D.G., M.H., J.K.A., W.L., T.L., Y.K.; Visualization H.G., M.H., and S.J.O.; Supervision, H.H.L., J.K.A., S.J.O., and W.L.; Funding Acquisition, H.H.L., J.K.A., and W.L.

## Declaration of Interests

Sarah J. Offutt and Yohan Kim are employees of Medtronic and own stock in the company. Jamu K. Alford was an employee and shareholder at Medtronic during the majority of this work.

## Methods

### Animal Surgical Preparations

The animal surgical procedures are detailed in previous work (Markovitz et al., 2013; Offutt et al., 2014; Straka et al., 2015). Experiments were performed on thirty young Hartley guinea pigs (350–580 g; Elm Hill Breeding Labs, Chelmsford, MA, USA) in accordance with policies of the University of Minnesota Institutional Animal Care and Use Committee. Each animal was anesthetized with an intramuscular injection of a mixture of ketamine (40 mg/kg) and xylazine (10 mg/kg) with supplements every 45–60 min to maintain an areflexive state. Heart rate and blood oxygenation was continuously monitored using a pulse oximeter (Edan Instruments Inc., Shenzhen, China), and body temperature was maintained at 38.0 ± 0.5°C using a heating blanket and rectal thermometer. Each animal was fixed in a stereotaxic frame (David Kopf Instruments, Tujunga, CA, USA), and a craniotomy was performed to expose the right auditory, visual and somatosensory cortices. Different configurations of 32-site electrode arrays (NeuroNexus Technologies, Ann Arbor, MI, USA) were inserted into the cortex and midbrain using hydraulic micro-manipulators (David Kopf Instruments, Tujunga, CA, USA). Example placements are shown in Figure 1A, 2C and 5A. After placement of the arrays, the brain was covered with agarose to reduce swelling, pulsations, and drying during recording sessions.

### Neural Recording and Stimulation

Experiments were performed within a sound attenuating, electrically shielded room using custom software and TDT hardware (Tucker-Davis Technology, Alachua, FL, USA). The electrode array used for A1 and SC1 consisted of four 5 mm-long shanks separated by 500 μm with eight iridium sites linearly spaced 200 μm (center-to-center) along each shank. The electrode array used for ICC in the auditory midbrain consisted of two 10 mm-long shanks separated by 500 μm with 16 iridium sites linearly spaced at 100 μm. The impedances were measured using niPOD (NeuroNexus Technologies, Ann Arbor, MI, USA) and ranged from 0.4–0.7 MΩ and 0.8–1.5 MΩ (at 1 kHz) for SC1/A1 and ICC probes, respectively. The A1 array was placed perpendicular to the cortical surface and inserted to a depth of approximately 1.6 mm such that the four shanks were arranged along the tonotopic gradient of A1. The ICC array was inserted 45° to the sagittal plane through the occipital cortex and into the ICC such that the sites spanned the tonotopic gradient of the ICC (Lim and Anderson, 2006; Snyder et al., 2004). Multi-unit neural data was recorded and sampled at a rate of 25 kHz, passed through analog DC-blocking and antialiasing filters up to 7.5 kHz, and digitally filtered between 0.3 and 3.0 kHz for the analysis of neural spikes; spikes were determined to be voltages exceeding 3.5 (or higher) times the standard deviation of the noise floor. All the PSTHs are plotted across 100 trials and binned at 1 msec, except for the PSTHs in Figure 6D,E which are binned at 2 msec for better visualization. All acoustic stimulation was presented to the animal’s left ear canal via a speaker coupled to a custom-made hollow ear bar. The speaker-ear bar system was calibrated using a 0.25” condenser microphone (ACO Pacific, Belmont, CA, USA). For guiding placement of the electrode array into the somatosensory cortex, electrical stimulation of the foot (biphasic, cathodic-leading pulse, 205 μs/phase, levels of 2.82 mA) with the stimulation ground placed in the foot was used. The input layer of A1 is layer III/IV in guinea pigs (Huang and Winer, 2000; Smith and Populin, 2001), which was identified by performing current source density (CSD) analysis as described in one previous study from our lab (Straka et al., 2015). The one-dimensional CSD approximation provides a consistent representation for the current sinks and sources associated with columnar synaptic activity in the guinea pig auditory cortex (Lim and Anderson, 2007; Middlebrooks, 2008). The main input layer of AC corresponded to the site with the shortest latency current sink (i.e., positive CSD peak). The layer II was identified as being two electrode sites (~400 μm) above the layer III/IV site.

### Verification of Probe Placement

Pure tones (50 ms duration, 5 ms ramp/decay) of varying frequencies (0.6–38 kHz, 8 steps/octave) and levels (0–70 dB SPL in 10 dB steps) were randomly presented to the animal’s left ear (4 trials/parameter), and acoustic-driven responses were recorded in A1 and the ICC to determine the functional location of each electrode site. A frequency response map was created for each recording site using driven spike rates (windowed 5–60 msec after tone onset for ICC sites and 5–20 msec for A1 sites) in which the spike rate was plotted on a color scale as a function of pure tone frequency (abscissa) and stimulus level (ordinate). From these frequency response maps, the best frequency for a given site was determined to be the frequency centroid at 10 dB above the visually determined neural threshold. For A1 placements, increasing best frequencies along the rostrolateral to caudomedial axis and short response latencies of approximately 15 msec confirmed that the array was within A1. Array placement within the ICC was confirmed by observing frequency response maps that increased in best frequency with increasing depth. In SC1 control experiments of US stimulation of visual cortex, the location of SC1 was confirmed by observing the evoked SC1 activity responding to electrical stimulation (biphasic, cathodic-leading pulse, 205 μs/phase, levels of 2.82 mA) of different body regions (e.g., foot, shoulder) as is expected for the somatotopic map of SC1 (Rapisarda et al., 1990).

### Electrode Site Reconstructions

In SC1 mapping experiments, site locations in SC1 were identified by imaging the exposed cortical surface with the inserted array shanks using a microscope mounted camera (OPMI 1 FR pro, Zeiss, Dublin, CA). Locations across animals were then normalized based on their relative distances from the middle suture line, bregma, and the lateral suture line, as successfully performed in previous studies from our lab (Markovitz et al., 2013, 2015b).

### Data Analysis

Response latencies for ICC sites and A1 sites were calculated using the first spike latency method. The time between the stimulus and the first spike following the stimulus was determined for each trial, and these times were averaged across all trials to determine the latency of each recording site. Spike counts for SC1 sites in response to US stimulation of visual cortex, electrical stimulation of the left foot, and electrical stimulation of the left shoulder were measured over a manually selected window 5–20 msec after the beginning of the onset of the stimulation to avoid electrical artifacts. Statistical comparisons between groups were performed using a ranked unequal variance two-tailed t-test (*P* < 0.05; Ruxton, 2006). The activated sites were determined by statistical testing if the evoked responses had higher spike rates than the spontaneous activity before the onset of stimulation (same window length as evoked responses) using a ranked unequal variance two-tailed t-test (*P* < 0.001).

### US Transducer Parameters and Placement

One US transducer at 220 kHz (Sonic Concepts, Bothell, WA) was used in this study, which was fitted with a 3D-printed focusing cone with a point diameter of 3 mm. This transducer was powered by a signal generator (Keysight Technologies 33512B, Santa Rosa, CA) amplified by a radio frequency amplifier (E&I 2200, Electronics & Innovation, Ltd., NY, USA). Through the signal generator, the primary channel was used to drive the US center frequency, and a secondary channel was used to modulate the center frequency signal into desired pulse envelopes. For A1 activation, the US transducer was placed over A1 and coupled directly to the brain via agarose gel. The A1 probe was placed such that electrical stimulation resulted in ICC activation, based on previous studies (Markovitz et al., 2015a). US paradigms (single pulse or 10 Hz-1.5 kHz PRF, 0.1–10 msec PD, 25 kPa-2 MPa pressure, 500 msec PD; Table S1) were tested in A1. For assessing SC1 activation, the probe was inserted into SC1 and the US transducer was placed over SC1 and coupled directly to the brain via agarose gel. US paradigms (single pulse or 1–16 kHz PRF, 0.31–10 msec PD, 50 kPa-1.6 MPa pressure, 1–6 sec PD; Table S3) were tested in SC1. In control experiments, the transducer was placed in several different locations, including directly on the brain with the skull and dura removed in A1, somatosensory cortex and visual cortex. For targeting the visual cortex with US, the same 32-site electrode array used in SC1 experiments was implanted into the lateral and caudal region of the right brain to first identify the visual cortex. The location of the visual cortex was confirmed when neural responses were elicited to light stimulation using white light-emitting diodes flashing at 2 Hz applied to the left eye.

### Ultrasound Calibration

Transducer outputs were characterized in a tank filled with deionized, degassed water under free-field conditions (i.e., without the presence of reflective obstacles). Each transducer/coupling cone unit was held perpendicularly above the water tank such that only the tip of the cone was submerged. A high sensitivity hydrophone (HNR 0500 ONDA Corp, Sunnyvale, CA, USA), calibrated between 50 kHz to 20 MHz, was positioned directly underneath the cone tip to acquire pressure measurements. Ultrasound output calibration charts were obtained based on peak negative pressure amplitudes collected at 1 mm below the tip of the coupling cone. Ultrasound field mapping was performed using a 3D motorized positioning system (Velmex Inc., Bloomfield, NY, USA). Pressure field profiles (Figure S1) were constructed by referencing negative pressure amplitudes of sonication pulses collected at different spatial locations, which were normalized with respect to the highest spatial peak negative pressure measured at the intended focus of the transducer.

### Additional Surgical Preparations for Control Experiments

For animals in which the auditory nerves were transected, additional surgical preparations were required. During the craniotomy, the skull was removed five millimeters rostral and caudal of the transverse cerebral fissure for the full width of the brain (i.e., as far lateral as possible before the skull descends ventrally). This step was performed prior to placing the ultrasound transducer or electrode arrays. After recordings were performed with the auditory nerves intact, the auditory nerves were cut using a fine needle which was inserted between the skull and the cortex at approximately 1–2 mm rostral of the transverse cerebral fissure and moved back and forth along the inner skull surface in a rostral-caudal direction to sever the auditory nerve. This step was performed on both sides of the head. For animals in which the cochlear fluids were removed, subcutaneous lidocaine was given to both ears and small incisions were made in the postauricular area to expose the skull overlying the bullas. The muscles around the incision were removed and a small hole was made to expose the bulla and provide access to the cochlea. A fine needle was inserted into the round window of the cochlea and the fluids were removed with a surgical vacuum drain (Schuco S130A, Allied Healthcare Products, Inc., St. Louis, MO). The cochlear fluid was removed for both ears. For animals connected with a gel channel, one 32-site electrode array was initially inserted into the A1 of the live animal then moved to the SC1 as described in animal surgical preparations section above. The brain of the live animal was then covered with a degassed-agar layer to prevent potential damage from direct contact with the ultrasound gel that connected the live animal with the skull of the dead animal. Transcranial US stimulation was presented to the skull of the dead animal (Figure 7E). Afterwards, the right and left side skull of the dead animal were both removed. A gel channel was built to connect the left exposed brain of the dead animal with the agar-covered right brain of the live animal. The US stimulation was then performed to the right brain of the dead animal without touching the gel channel (Figure 7H).

